# Taxonomic and functional anuran beta diversity of a subtropical metacommunity respond differentially to environmental and spatial predictors

**DOI:** 10.1101/588947

**Authors:** Diego Anderson Dalmolin, Alexandro Marques Tozetti, Maria João Ramos Pereira

## Abstract

The relative contributions of environmental and spatial predictors in the patterns of taxonomic and functional anuran beta diversity were examined in 33 ponds of a metacommunity along the coast of south Brazil. Anurans exhibit limited dispersion ability and have physiological and behavioural characteristics that narrow their relationships with both environmental and spatial predictors. So, we expected that neutral processes and, in particular, niche-based processes could have similar influence on the taxonomic and functional beta diversity patterns. Variation partitioning and distance-based methods (db-RDA) were conducted with presence/absence and abundance data to examine taxonomic and functional facets and components (total, turnover and nestedness-resultant) in relation to environmental and spatial predictors. Processes determining metacommunity structure were similar between the components of beta diversity but differed among taxonomic and functional diversity. While taxonomic beta diversity was further accounted by environmental predictors, functional beta diversity responded more strongly to spatial predictors. These patterns were more evident when assessed through abundance data. These opposing patterns were contrary to what we had predicted, suggesting that while there is a taxonomic turnover mediated by environmental filters, the spatial distance promotes the trait dissimilarity between sites. Our results reinforce the idea that studies aiming to evaluate the patterns of structure in metacommunities should include different facets of diversity so that better interpretations can be achieved.

## 1 INTRODUCTION

Beta diversity connects the spatial structure of communities to a variety of ecological processes, such as neutral processes (e.g. dispersion limitation) and niche-based processes (e.g. limiting similarity and environmental filtering) [1, 2, 3, 4]. Beta diversity represents the amount of variation in species composition between a set of local communities 2, 5, 6], and can be divided into two distinct components: spatial turnover and nestedness [7, 8]. Spatial turnover occurs when some species are replaced by others as a result of ecological processes (e.g. environmental filtering and dispersion limitation) that restrict their occurrence in certain places [2, 8, 9]. Nestedness, on the other hand, occurs when non-random processes of species loss result in the ordered deconstruction of assemblages, leading to the formation of local sets poorer in the number of species, and subsets of richer sites [9, 10, 11]. Despite their distinct nature, these two components are complementary and main drivers of dissimilarity patterns between communities [7].

Within the metacommunity theory, the organization of local assemblies is thought to occur at broader spatial scales [12, 13]. In metacommunities structured by neutral processes, the dispersion limitation and demographic stochasticity are the dominant factors, so that geographical distance between communities is the best predictor of beta diversity [14, 15]. In contrast, ecological interactions and environmental conditions are the most important factors in metacommunities structured by niche processes, and the environmental distance between communities should be the best predictor of beta diversity [12, 16]. However, several studies suggest that the action of these processes is not mutually exclusive, and that the structuring of biological metacommunities results from the interaction of the two processes – neutral and niche-based [13, 17]. Indeed, in aquatic metacommunities deterministic processes (especially environmental filtering) seem to be dominant, though neutral processes also contribute to the observed patterns of beta diversity [9].

Taking into account that biological communities result from a complexity of interactions between organisms, environment and space, the incorporation of functional traits (functional diversity) in community and metacommunity studies has been widely advocated (e.g. 18, 19). In fact, the use of an integrative approach where taxonomic and functional diversities are taken into account is advantageous. First, the evaluation of communities using only taxonomic identity is often difficult to interpret, since taxonomic groups may contain phylogenetic and ecological lineages in conflict between convergence and adaptive divergence [20, 21, 22]. In addition, the functional approach allows elucidating the ‘true role’ of each species in ecosystem processes and their resistance and resilience to environmental changes [23, 24]. Finally, several studies found congruent responses of the two metrics of diversity for the same ecosystem processes [19, 25, 26], although these relationships may vary according to the taxonomic group of interest [27, 28].

Neotropical anurans are considered excellent ecological models because they are locally abundant and sampling of most groups is relatively easy. Anurans have highly permeable skin, a complex and biphasic life cycle, limited dispersion, and geographically restricted distribution patterns [29, 30]. So, compositional variation in anuran seems to result from several factors, such as available area and hydroperiod, vegetation cover, type of surrounding matrix and geomorphology [26, 31, 32, 33, 34]. Thus, both environment and space tend to strongly contribute to the patterns of taxonomic and functional dissimilarity between anuran communities [30, 35, 36].

Although anuran beta diversity has already been addressed in several studies (e.g. 33, 36, 37), few have used an integrative approach to describe patterns of anuran beta diversity. In this study we investigated the relationship between environmental and spatial components and the patterns of taxonomic and functional beta diversity in a metacommunity of anurans from the coastal subtropical region of southern Brazil. We expect beta diversity to be influenced by both environmental and spatial components, with a greater contribution of the environmental component to the distribution of traits and species [38]. We also expect diversity components – taxonomic and functional – to present similar responses to the sets of descriptors evaluated [30]. Through functional traits, species can shape, change and accommodate in the environment where they occur [39, 40]. Consequently, species distribution can be expected as resulting from combinations of ecologically relevant characteristics allowing them to persist in a given set of environments [41]. We thus expect functional diversity to be a better indicator of the ecological processes responsible for the structuring of the anuran metacommunity [38, 42].

## 2 Material and Methods

### 2.1 Ethics statement

Collection permits were provided by Instituto Chico Mendes de Conservação da Biodiversidade (ICMBio) (authorization 55409). Field studies did not involve endangered or protected species. We restricted manipulation of animals in the field to minimal as we sampled just specimes restricted in the collection units (see the section 2.3). Specimens collected were identified, measured and immediately released after these procedures in the same pond where they were sampled. All sampling procedures were reviewed and specifically approved as part of obtaining the field permits by ICMBio (see above).

### 2.2 Study Area

This study was done in Lagoa do Peixe National Park (PNLP; 31°02_- 31°48_S; 50°77_-51°15_W; figure 1), the only Ramsar site in southern Brazil [43]. With a length of 64 km and an average width of 6 km, the PNLP comprises over 34,000 hectares of protected wetlands, integrating the Coastal Plain of the State of Rio Grande do Sul, one of the regions of southern Brazil with higher concentration of wetlands [44]. The climate is subtropical humid, and temperatures range between 13 °C and 24 °C with annual average of 17.5 °C. The mean annual precipitation varies between 1200 and 1500 mm [45].

**Figure 1:**
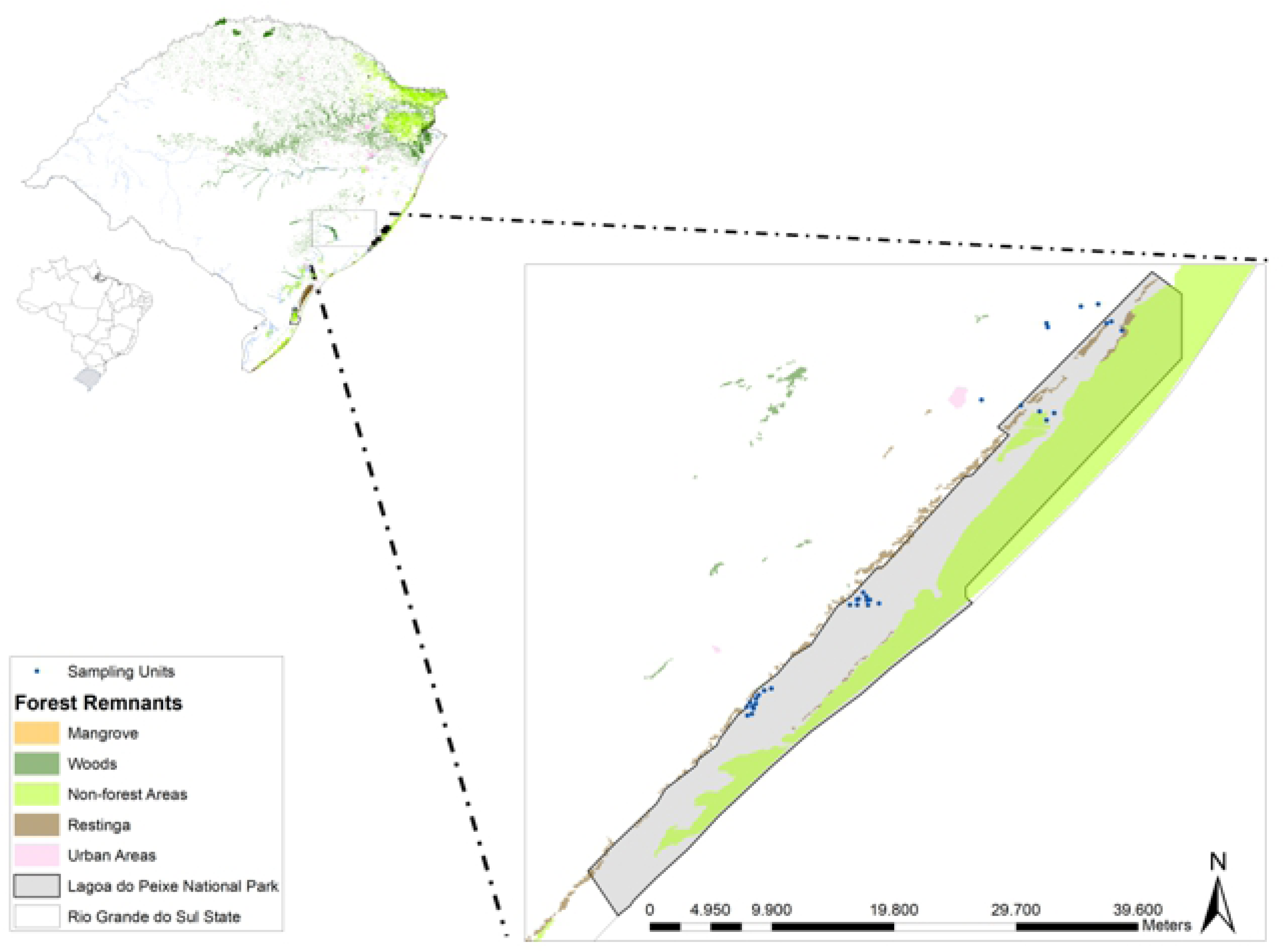
Study area at Lagoa do Peixe National Park, southern Brazil. Sampled ponds are represented by blue circles (N = 33).

### 2.3 Anuran surveys and trait measurement

We sampled adult anurans in 33 water bodies throughout the study region (Fig. 1). Ponds were selected based on biotic and abiotic characteristics (see Table S2), insulation of other ponds (sample independence), accessibility and landowner permission. Distances between ponds ranged from 0.7 to 39 km. Sampling was performed monthly, from October 2016 to March 2017. We used both calling surveys and active search at breeding habitats to record the number of calling males of each species in each pond [46]. Samplings were done from 6 p.m. to 0 a.m. The total effort per pond was 1 hour per month, totalling 6 hours of sampling per pond.

Five morphological traits we meansuared for each individual captured: head shape; eyes position; eye size; relative length of limbs; body mass (see Supplementary Material Table S1 for more information). Addittionaly, we compiled six life history traits from literature: reproductive mode; relative number of eggs; daily activity period; type of habitat; fossorial habit; reproductive season. These 11 traits were chosed based on perceived importance for determining habitat use and species resilience. All attributes were used to construct a pairwise distance matrix of species. In this procedure we used the Gower standardization for mixed variables [47].

### 2.4 Environmental and spatial variables

The environmental descriptors measured in this study and their description are presented in Table S2. Area, depth and number of vegetation types around the pond, pond vegetation (inside the pond), and margin configuration and pound substrate were measured in the field through visual interpretation in an area around 5 meters from the edge of the ponds. The distance to the nearest forest fragment and distance to the nearest sampled pond were obtained from high-resolution aerial photographs of the region inspected immediately after samplings, available from Google Earth (http://earth.google.com/), combined with field inspection.

We used distance-based Moran’s Eigenvector Maps (dbMEMs; 47, 48) to create spatial variables (eigenvectors) based on the Euclidean distance matrix of the geographical coordinates of the ponds. First of all, we defined the neighbourhood matrix, which describes the spatial relationships among objects [48]. In other words, we defined which ponds are neighbours and which are not. We used as spatial neighbourhood graphs ‘Delaunay triangulation’, ‘Gabriel graph’ and ‘Minimum spanning trees’, and as a weight measure we used the linear distances between ponds. We selected the best neighbourhood matrix based on AICc. The most parsimonious model was the one based on the ‘Gabriel graph’, and the truncation distance was 18.03 km (AIC = 11.22 versus 12.88 for the null model). This model also generated 23 spatial variables (eigenvectors), eight of which with positive autocorrelation.

### 2.5 Data Analysis

#### 2.5.1 Data matrices

We built four matrix types containing the measured data of all ponds: (i) an abundance and/or presence/absence matrix, which contained the total species count for each pond; (ii) a trait matrix, containing average trait values foreach species; (iii) an environmental matrix, containing all environmental descriptors measured in each pond; and (iv) a spatial matrix, containing all dbMEMs.

As suggested by [49], we considered the abundance of each species in a given pond as the abundance of calling males recorded in the month of highest recorded abundance. This procedure prevents underestimates of population abundance caused by calculating the mean of successive samples and prevents overestimates caused by the re-counting of individuals if successive samples are summed [49]. Abundance data were transformed using the Hellinger distance [47] to homogenize variation among species abundances. Environmental descriptors values were standardized by subtracting each value from the average of the descriptor and dividing the result by the standard deviation.

#### 2.5.2 Assessing the anuran beta diversity components

Following the methods proposed by [8], we partitioned beta diversity into overall beta diversity, turnover and nestedness components. This procedure was performed using the function “beta.pair” in the R package betapart [50] and the function “beta” in the R package BAT [51]. We used the Sorensen dissimilarity for the presence/absence data and the Bray-Curtis dissimilarity for the abundance data. This procedure produces three dissimilarity matrices (Table 1).

**Table 1:** Summary of beta diversity index and their nomenclatures used in this study

| Beta diversity index |  | Type of data |  |
| --- | --- | --- | --- |
|  |  | Presence/absence | Abundance |
| Total | Overall spatial turnover | $\beta_{\text{Sor}}$ | $\beta_{\text{Bray}}$ |
| Turnover | Turnover immune to species richness variation | $\beta_{\text{Sim}}$ | $\beta_{\text{Bal}}$ |
| Nestedness | Nestedness resulting from species richness differences between sites | $\beta_{\text{Sne}}$ | $\beta_{\text{Gra}}$ |

#### 2.5.3 Community–environment relationships

We used distance-based redundancy analysis (db-RDA) on each biological dissimilarity matrix to examine community–environment relationships in more detail [52]. This method is similar to redundancy analysis, but may be based on any dissimilarity or distance matrix (in our case, Sorensen and Bray-Curtis dissimilarities) [47].

Initially we selected only significant environmental predictors of variation in taxonomic and functional beta diversity, which then were used for the environmental model [53]. We used the bio-env analysis [54] to produce ‘the best’ environmental distance matrix. This method is based on standardized environmental variables and tests all possible combinations of the environmental variables, providing information as to which combination shows the strongest correlation between biological dissimilarity and the environmental distance matrix. The bio-env analyses were based on the Pearson correlation coefficient and 9999 permutations for obtaining *p*-values. All environmental descriptors were included (continuous and categorical). We ran these tests using the functions “bioenv,” from the R package vegan [55]. The tables with bioenv selection procedure results are presented in the Tables S3 and S4.

#### 2.5.4 Spatial predictors

We used forward selection with 9999 permutations to select the MEMs to run the spatial model. The selection stopped either when the tested variable had a *p*-value above 0.05 or when the adjusted *R*^*^2^*^ [56] of the full model, before any selection, was exceeded [53]. The forward selection procedure was run with the “forward.sel” function from the R package vegan [53]. The summary results of forward selection procedure are presented in the Tables S5 and S6.

#### 2.5.5 Variation partitioning for the anuran taxonomic and functional beta diversity

The relative contributions of the environmental descriptors and spatial variables to the taxonomic and functional beta diversity patterns were evaluated using a partial Redundancy Analysis (pRDA) with variation partitioning [48]. This analysis partitions the variance in community composition resulting from each explanatory variable ([E] = environment and [S] = spatial), (2) the unique contribution of each explanatory variable ([E/S] = environment - purely environmental variables – or [S/E] = spatial – purely spatial variables) and (3) the total variance explained by the environmental and spatial variables together (spatially structured environmental variables). The variance explained by each fraction was based on the adjusted *R*^2^ [56]. The environmental variables used in the environmental model and the spatial variables composing the spatial model were those previously selected in db-RDA, bioenv selection and forward selection, as described above.

The significance of db-RDA axes (Tables S7 and S8) and pRDA fractions were tested through an ANOVA-like permutation test to assess the significance of the constraints, using 9999 permutations. The db-RDA and pRDA analyses were done using the functions “capscale” and “var.part”, and the permutations using the “anova.cca” function, of the R package vegan [55].

## 3 Results

During the sampling period, 11 species belonging to three families (Bufonidae, Hylidae and Leptodactylidae) were registered. The most frequent species were *Dendropsophus sanborni* and *Pseudis minuta*, occurringin 19 of 33 ponds evaluated. *Physalaemus biligonigerus* and *Scinax fuscovarius* were less frequent occuring, respectively, in three and four of the sampled ponds (for the complete list of species and occurrence pattern across ponds see Table S9).

### 3.1 Environmental predictors and anuran beta diversity

The “bioenv” function identified the most significant variables explaining the variation on taxonomic and functional beta diversity. In general, the sets of descriptors selected to compose the environmental and spatial models were similar between the presence/absence and the abundance data for total beta diversity and its components (turnover and nestedness; Tables 2 and 3; Tables S3 and S4). However, these sets were different between the taxonomic and the functional beta diversities. The greater relationship between environmental descriptors and taxonomic and functional beta diversity was observed in the total beta diversity, especially with abundance data (respectively: *R*^2^*adj*: 0.18; *F*: 1.52; *p*: <0.001; *R*^2^*adj*: 0.06; *F*: 1.99; *p*: 0.04).

**Table 2.**
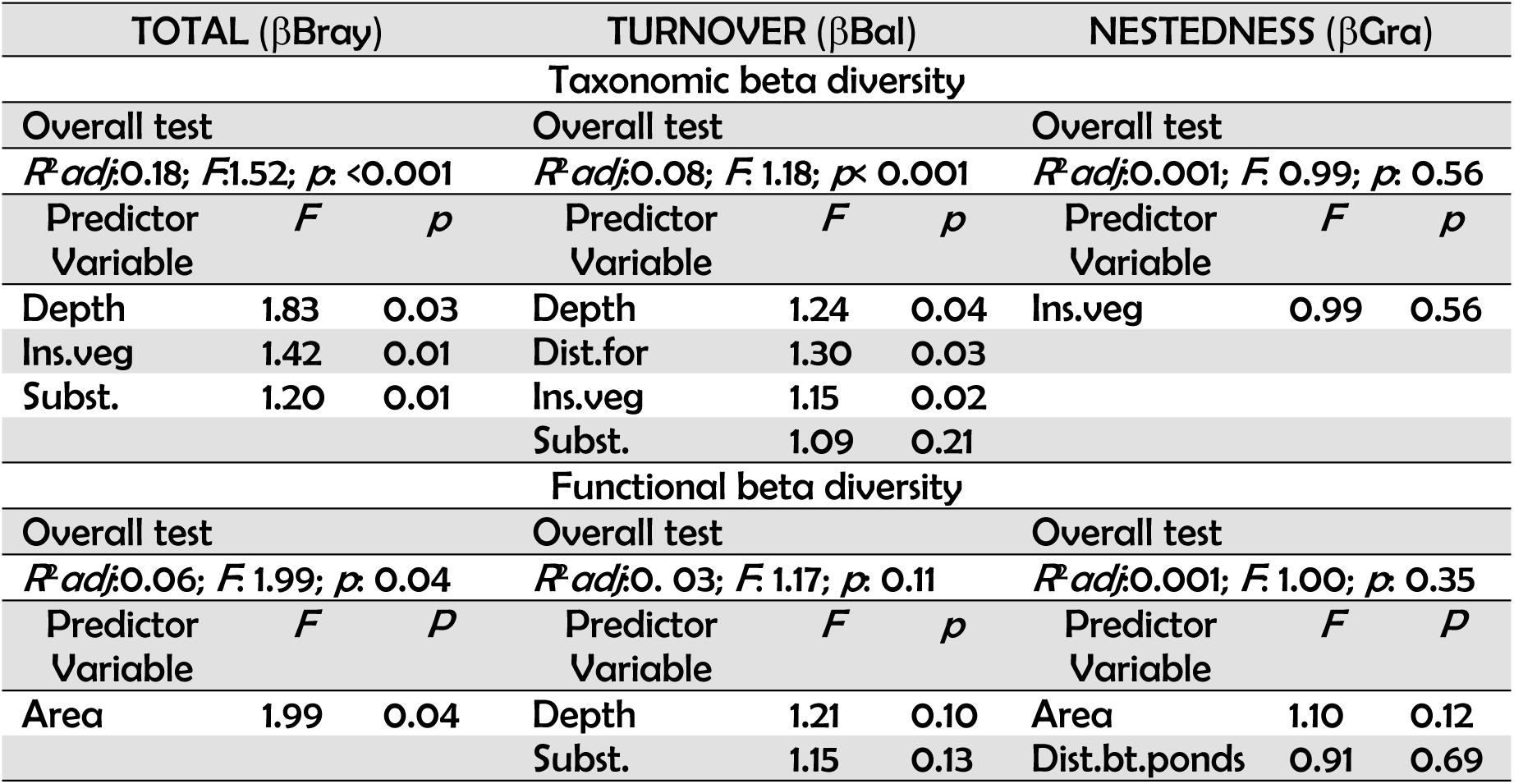
Results of distance-based RDAs for the abundance data. Analyses were run for taxonomic and functional components based on Sorensen, Simpson and nestedness-resultant dissimilarities. Full models and marginal tests of significance for single environmental variables are shown (i.e., separate significance test for each variable in a model when all other terms are in the model).

**Table 3.**
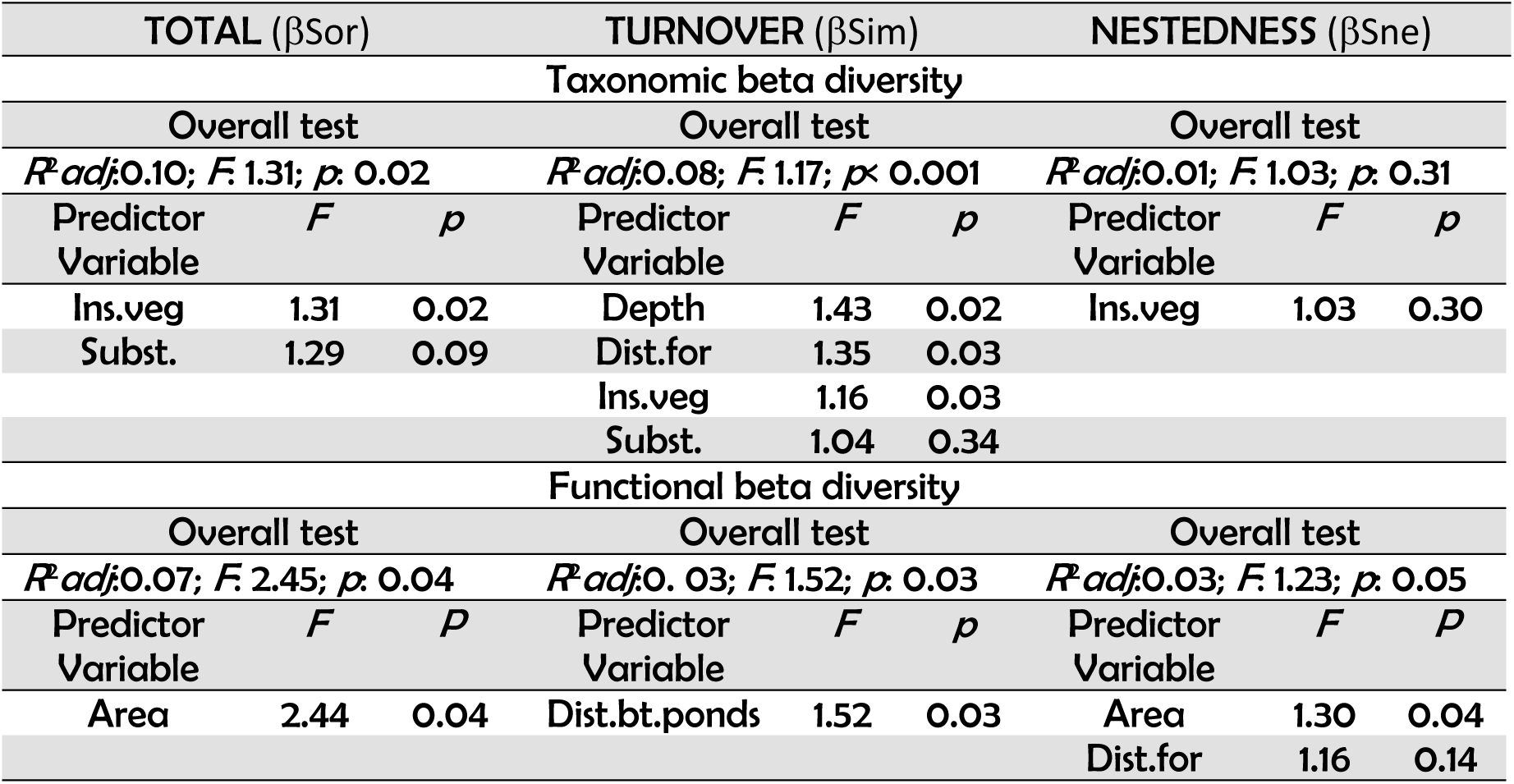
Results of distance-based RDAs for presence/absence data. Analyses were run for taxonomic and functional components based on Sorensen, Simpson and nestedness-resultant dissimilarities Full models and marginal tests of significance for single environmental variables are shown (i.e., separate significance test for each variable in a model when all other terms are in the model).

For taxonomic beta diversity, the environmental descriptors selected were depth, distance to the nearest forest, pond vegetation and pond substrate. Pond vegetation was shared for all of beta diversity components, but it was not significant for the nestedness component. Depth seems to be one of the main drivers of the taxonomic beta diversity in the metacommunity, since it largely explained total beta diversity (**βBray**: *F*=1.83, *p*=0.03) and turnover (**βBal**: *F*=1.24, *p*=0.04; **βSim**: *F*=1.43, *p*=0.02).

The environmental descriptors selected for the functional beta diversity were area, depth, distance between ponds, distance to the nearest forest and types of substrate. The sets of environmental descriptors selected were similar when using abundance or presence/absence data. Area was the descriptor that most explained total beta diversity (**βBray**: *F*=1.99, *p*=0.04; **βSor**: *F*=2.44, *p*=0.04) and nestedness (**βSne**: *F*=1.30, *p*=0.04). For the abundance data, dbRDA did not find any relationship between environmental descriptors, turnover and nestedness (**βBal**: *R*^2^*adj*=0. 03; *F*= 1.17; *p*= 0.11; **βGra**: *R*^2^*adj*=0.001; *F*= 1.00; *p*= 0.35).

#### 3.2 Variation partitioning for anuran beta diversity

Variation partitioning analyses identified significant effects of both environmental and spatial components on the anuran metacommunity structure. However, the relationship of these components was discrepant between the taxonomic and the functional beta diversities. While environment mostly influenced the taxonomic structure, the functional structure was mostly determined by the spatial component (Figure 2a-2d; Tables 4 and 5). Also, the relationship between beta diversity and environment and space was better explained for the abundance data.

**Table 4:** Variation partitioning for the components of anuran beta diversity based on **abundance** data. The table shows the variation explained (*R*^*^2^*^ *adjusted*) for Model 1 (turnover versus environment and space) and Model 2 (nestedness versus environment and space). E = environment; S = spatial component obtained from dbMEM; E+S = shared contribution between environment and space; E/S = the unique contribution of the environmental component; S/E = the unique contribution of the spatial component.

| | | Model 1 = TURNOVER ($\beta_{Bal}$) | | | Model 2 = NESTEDNESS ($\beta_{Gra}$) | | |
| --- | --- | --- | --- | --- | --- | --- | --- |
| | | $R^2$adjusted | $F$ | $p$ | $R^2$ adjusted | $F$ | $p$ |
| **FUNCTIONAL**<br>**BETA**<br>**DIVERSITY** | E | 0.21 | 2.62 | 0.05 | 0.13 | 2.36 | 0.09 |
|  | S | 0.79 | 9.67 | <0.001 | 0.71 | 9.69 | <0.001 |
|  | E+S | 0.24 | 6.35 | <0.001 | 0.11 | 8.10 | <0.001 |
|  | E/S | <0.01 | 0.60 | 0.76 | 0.03 | 1.67 | 0.18 |
|  | S/E | 0.56 | 5.43 | 0.002 | 0.60 | 8.25 | <0.001 |
|  | Residuals | 0.23 | - | - | 0.27 | - | - |
| **TAXONOMIC**<br>**BETA**<br>**DIVERSITY** | E | 0.31 | 2.01 | <0.001 | 0.06 | 1.23 | 0.27 |
|  | S | 0.34 | 2.77 | <0.001 | 0.24 | 3.50 | 0.003 |
|  | E+S | 0.26 | 1.87 | 0.009 | <0.01 | 2.26 | 0.004 |
|  | E/S | 0.06 | 1.14 | 0.33 | 0.11 | 1.49 | 0.05 |
|  | S/E | 0.08 | 1.25 | 0.29 | 0.28 | 3.44 | 0.003 |
|  | Residuals | 0.60 | - | - | 0.60 | - | - |

**Table 5:** Variation partitioning for the components of anuran beta diversity based on **presence/absence** data. The table shows the variation explained (*R*^*^2^*^ *adjusted*) for Model 1 (turnover versus environment and space) and Model 2 (nestedness versus environment and space). E = environment; S = spatial component obtained from dbMEM; E+S = shared contribution between environment and space; E/S = the unique contribution of the environmental component; S/E = the unique contribution of the spatial component.

| | | Model 1 = TURNOVER ($\beta_{Sim}$) | | | Model 2 = NESTEDNESS ($\beta_{Sne}$) | | |
| --- | --- | --- | --- | --- | --- | --- | --- |
| | | $R^2$ adjusted | $F$ | $p$ | $R^2$ adjusted | $F$ | $p$ |
| **FUNCTIONAL BETA DIVERSITY** | E | 0.20 | 3.24 | 0.05 | 0.17 | 2.85 | 0.03 |
|  | S | 0.41 | 5.20 | 0.004 | 0.05 | 1.45 | 0.18 |
|  | E+S | 0.24 | 3.14 | 0.02 | 0.06 | 1.84 | 0.05 |
|  | E/S | <0.01 | 0.53 | 0.84 | 0.11 | 2.04 | 0.05 |
|  | S/E | 0.17 | 2.47 | 0.03 | <0.01 | 0.88 | 0.51 |
|  | Residuals | 0.62 | - | - | 0.84 | - | - |
| **TAXONOMIC BETA DIVERSITY** | E | 0.34 | 2.13 | 0.002 | 0.11 | 1.43 | 0.14 |
|  | S | 0.11 | 2.48 | 0.007 | 0.19 | 3.49 | 0.01 |
|  | E+S | <0.01 | 2.63 | 0.003 | 0.08 | 1.71 | 0.04 |
|  | E/S | 0.36 | 1.35 | <0.001 | 0.10 | 1.40 | 0.15 |
|  | S/E | 0.13 | 2.56 | 0.009 | 0.08 | 2.66 | 0.02 |
|  | Residuals | 0.51 | - | - | 0.82 | - | - |

**Figure 2:**
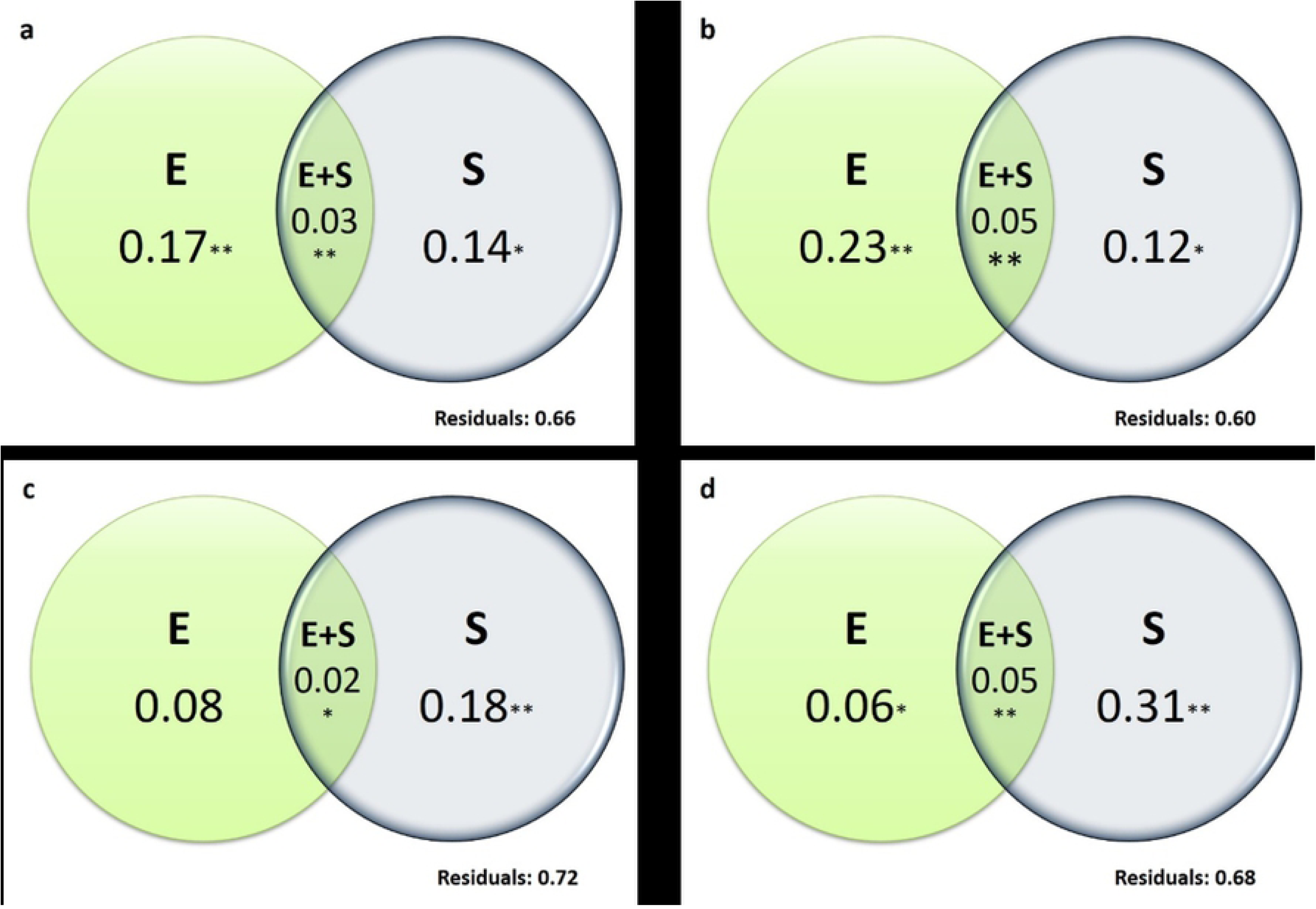
Variation partitioning for total taxonomic (a and b) and functional (c and d) beta diversities of anuran communities. Figures 2a and 2c: presence/absence data; Figures 2b and 2d: abundance data; E = environment; S = spatial component. (*) *p*< 0.05; (**) *p*<0.001.

#### 3.2.1 Variation partitioning for taxonomic beta diversity

The contribution of the environmental component for the total variance explained by the taxonomic beta diversity and its components was greater than the spatial component (Figure 2a and 2b, tables 4 and 5). For total beta diversity, the contributions of the environmental individual fraction were 17% (**βSor**: *F*=1.59, *p*=0.02) and 23% (**βBray**: *F*=1.79, *p*=0.004), while the contributions of space were 14% (**βSor**: *F*=1.98, *p*=0.04) and 12% (**βBray:** 6%; F=1.92, *p*=0.04). The shared contribution between Environment and Space on the total variance was 3% (**βSor**: *F*=1.94, *p*=0.003) and 5% (**βBray**: *F*=2.21, *p*<0.001).

The components of taxonomic beta diversity showed more complex patterns. The environmental, but mostly the spatial component, explained the nestedness patterns for the abundance data (**βGra**= 11% and 28%, respectively; Table 4). The shared fraction between the two components, environmental and spatial, explained < 1% of the variation (though it was significant; p=0.004). The opposite pattern was observed for the turnover, where only the shared fraction between environment and space was significant (**βBal**= 26%). For the presence/absence data, the environmental and spatial individual fractions were only significant in explaining species turnover, accumulating 36% and 13%, respectively (Table 5). The shared fraction between the two components, however, explained < 1% of the turnover variation. The environmental fraction was slightly larger that the spatial fraction in explaining nestedness (**βSne**= 10% and 8%, respectively), though the environmental fraction was not significant but the spatial fraction was (Table 5). The shared contribution between environment and space explaining nestedness was larger than that explaining the turnover (**βSne**= 8%).

#### 3.2.2 Variation partitioning for functional beta diversity

Variance in functional beta diversity and its components were mostly explained by spatial processes for both presence/absence (S: 18%; E: 8%) and abundance data (S: 31%; E: 6%) (Figure 2c and 2d, tables 4 and 5). The individual fraction of the environmental component was only significant for the abundance data (**βBray:** F=2.52, *p*=0.05). The shared contribution between environment and space for the total functional beta diversity variance was 2% (**βBray**: *F*=2.21, *p*<0.001) and 5% (**βSor**: *F*=5.18, *p*=0.005), respectively.

Components of functional beta diversity followed the same pattern as that of total beta diversity. For the abundance data there were significant contributions of the spatial component (**βBal**= 56%; **βGra**= 60%; Table 4) and of the shared fraction between space and environment (**βBal**= 24%; **βGra**= 11%). For the presence/absence data there were some discrepant results between turnover and nestedness. While the individual fraction of space significantly explained the turnover (**βSim**= 17%), the environmental individual fraction significantly explained nestedness (**βSne**= 11%). The shared fraction of space and environment were significant for both turnover and nestedness, but contributing mostly towards the turnover (**βSim**= 24%; **βSne**= 6%).

## 4 DISCUSSION

Here, we used a variation partition approach to understand how taxonomic and functional anuran beta diversity are influenced by environment and space at a regional scale in South American subtropical wetlands. Our results showed opposing patterns between taxonomic and functional beta diversity in their response to environmental and spatial predictors, contrary to what we had predicted, and has been described for metacommunities of Atlantic Forest anurans [30]. Taxonomic diversity responded mostly to local environmental predictors, while functional diversity is better explained by spatial predictors. These patterns were evident when we used both presence/absence and abundance data, though, abundance data evidenced those patterns more effectively, as reported in other studies [57, 58].

Our findings suggest thus that the structuring of beta diversity in metacommunities of organisms that depend on aquatic systems, including those of anurans, is complex and responds differently to environmental and spatial predictors, and this is in agreement with the results of studies using similar approaches [22, 30]. This allows us to infer that different processes act on the selection of species and functional attributes along metacommunities [22].

Taxonomic and functional nestedness were explained almost exclusively by purely spatial predictors. Turnover, however, presented complex patterns similar to those observed for total beta diversity. These results are in agreement with those of [8], who initially demonstrated that the underlying patterns and processes of diversity might differ when total beta diversity is partitioned. Our results also support the assumption that beta diversity components vary across geographic space and may respond to different predictors [1, 12].

### 4.1 Environmental predictors and anuran beta diversity

The organization of anuran assemblages in freshwater systems exhibits patterns in response to environmental gradients [29, 34, 59]. Most of the descriptors we evaluated had some level of influence on anuran patterns of beta diversity. Indeed, different environmental factors tend to affect species differently due to differences in their physiological and behavioural characteristics [20, 35, 60. Beta diversity was mostly driven by area and depth. These two variables are associated with pond hydroperiod and promote a trade-off between the persistence of those systems and predation and competition levels, both with strong influence in anuran survival and life cycle [31, 32]. Species richness and composition will thus be affected by those descriptors, and the persistence of a species in a given community will be mediated by the presence of specific traits [31, 32]. For example, species that lay eggs in foam nests and show rapid larval development (Leptodactylidae e.g. *Physalaemus biligonigerus* and *Physalaemus gracilis*) survive in shallow and drought-prone areas but tend to present reduced rates of growth and post-metamorphic survival [61, 62]. On the other hand, the occurrence in permanent ponds is favoured by morphological and behavioural traits that facilitate the escape and co-occurrence with predators and competitors, respectively (e.g. body and fin format; refuge use; activity patterns) at the cost of time delays in development and phenotypic changes promoted by intra and interspecific interactions [63].

Substrate type and pond and margin vegetation are closely associated with species’ reproductive habits (e.g. calling sites and oviposition sites). Anurans present a wide variety of reproductive modes [64, 65], and the presence of more differentiated modes requires increasing levels of environmental complexity. This is the case for most species of the Hylidae and the Leptodactylidae, which place spawning near aquatic macrophytes [65]. Distance between ponds and to the nearest forest fragment may be spatially structured and associated with the dispersion of individuals [29, 66]. Indeed, other ponds and closely-located forest patches may function as a temporary refuge or permanent site (as the case of species that show high site fidelity) [67] for feeding and thermoregulation [68, 69], and seem to be essential for maintaining richness and abundance of anurans in local communities across a variety of regions (e.g. Europe - [70]; North America - [32]; South America - [66]).

### 4.2 Opposing patterns between taxonomic and functional beta diversity

Our results show that taxonomic beta diversity is mainly structured by niche-based processes, as the similarity in species composition decays along the environmental gradient [12]. This means that environmentally distinct ponds add different community compositions [71, 72]. However, this dissimilarity was lower for the functional diversity, indicating that, although there is a taxonomic turnover mediated by environmental filters, the traits present along these gradients are not sufficiently different for large functional dissimilarities to be observed [25, 73].

Functional beta diversity was more closely associated with spatial predictors, suggesting the dominance of neutral processes [14]. The presence of a spatial structure in beta diversity patterns of anurans has been observed to communities of Amazonia [66] and southeastern Brazil [29, 60] and appears to be common and more evident for amphibians than for other organisms (e.g. mammals, birds and invertebrates)[1]. We highlight that the combination of spatial and environmental models also contributed considerably to the explained fraction, which may suggest a certain level of spatial autocorrelation in some of the environmental variables [3].

The results for taxonomic nestedness and functional nestedness and turnover emphasize the pattern of dominance by spatial predictors in our study area. This may be linked to dispersal limitation (e.g. [36]); indeed, anurans relatively small body-size and physiological limitations make most species dependent on flooded or at least humid corridors for dispersal [20, 32]. In addition, the presence of natural and artificial barriers (e.g. sandy soils and roads with constant car traffic, as occurs in our study area) may prevent many species from reaching all ponds available for the whole of the metacommunity [1, 32]. As a consequence, there was a decrease in functional similarity with the increase in geographical distance, as occurs in other aquatic organisms [22, 74, 75].

Still, the association of spatial predictors with neutral dynamics and potential patterns of dispersal limitation should be interpreted cautiously. In fact, potentially important environmental variables may not have been evaluated, while contained in the spatial component [72, 76]. Also, other factors difficult to evaluate, such as predation and competition, may have a significant effect in the patterns of beta diversity and alter the importance of each predictor along the metacommunity, which was already demonstrated for microorganisms in controlled systems [77].

The processes of community assembly may act in different environmental gradients and along spatial and temporal scales, resulting in patterns of functional and phylogenetic convergence or divergence independent of the taxonomic identity [27, 78]. Such discrepancies between different metrics of diversity across spatial and temporal scales were already reported in other taxa [19, 58, 79]. Moreover, the correlation between taxonomic and functional nestedness and turnover may be higher at the alpha scale than at the beta and may be variable according to the set of the functional traits evaluated [3, 80, 81].

## 5 CONCLUSIONS

In conclusion, our results contributed to the knowledge about the relative influence of neutral and niche-based processes in determining the structure of anuran metacommunities. Along the southern coast of Brazil, the structure of beta diversity differed between the taxonomic and functional components. Despite this, we found important congruences between the components of beta diversity within each facet. The dominance of environmental predictors in the structuring of the taxonomic diversity and the spatial predictors in the structuring of the functional diversity suggest the co-existence of different processes in structuring anuran metacommunities and reinforce the importance of the inclusion of different facets of diversity in such analyses. The substantial contribution of purely spatial predictors in the patterns of both facets of diversity is similar to that found in other regions (e.g. [30]), confirming the predominance of niche-based processes on the structuring of anuran metacommunities. However, the significant contribution of the shared fraction between environment and space, shows that the structure of our target metacommunity results from the interaction of both processes [17].

## 6 ACKNOWLEDGMENTS

We thank Conselho Nacional de Desenvolvimento Científico e Tecnológico (CNPq) for financial and logistical support and for the Diego Dalmolin’ Phd scholarship and Maria João Pereira’ CNPq Research Productivity scholarship. We also thank Lagoa do Peixe National Park for all the support during the field activities. This study was done under ICMBIO licence number 55409-1. Finally, we thank all those who helped during field and lab work and the Márcio Borges Martins (UFRGS) and Vinícius Bastazini (CNRS) for their contributions to previous versions of this manuscript.

## 7 Supporting information captions

**Table S1 -** Functional traits measured (in adults). (DOCX)

**Table S2–** Environmental descriptors of ponds measured between October 2016 and March 2017 at Lagoa do Peixe Nation Park, Rio Grande do Sul, Brazil. (DOCX)

**Table S3** – Results of bioenv selection for the environmental variables selected to compose the environmental model of taxonomic beta diversity during the db-RDA and pRDA analysis. (DOCX)

**Table S4** – Results of bioenv selection for the environmental variables selected to compose the environmental model of functional beta diversity during the db-RDA and pRDA analysis. (DOCX)

**Table S5** – Results for the forward selection of spatial variables to compose the spatial model of taxonomic beta diversity during the pRDA analysis. (DOCX)

**Table S6** – Results of forward selection of spatial variables to compose the spatial model of functional beta diversity during the pRDA analysis. (DOCX)

**Table S7**– Results of anova.cca test for the two first axis of db-RDA between environmental variables selected and functional beta diversity components. (DOCX)

**Table S8** – Results of anova.cca test for the two first axis of db-RDA between environmental variables selected and taxonomic beta diversity components. (DOCX)

**Table S9 –** Anuran species composition in each of the sampled ponds. (DOCX)

## 8 Author Contributions

Conceptualization: Diego Anderson Dalmolin, Maria João Ramos Pereira, Alexandro Marques Tozetti.

Data curation: Diego Anderson Dalmolin.

Formal analysis: Diego Anderson Dalmolin, Maria João Ramos Pereira.

Funding acquisition: Diego Anderson Dalmolin, Maria João Ramos Pereira.

Investigation: Diego Anderson Dalmolin, Maria João Ramos Pereira, Alexandro Marques Tozetti.

Methodology: Diego Anderson Dalmolin, Maria João Ramos Pereira, Alexandro Marques Tozetti.

Project administration: Maria João Ramos Pereira.

Resources: Diego Anderson Dalmolin, Maria João Ramos Pereira.

Supervision: Diego Anderson Dalmolin, Maria João Ramos Pereira.

Writing ± original draft: Diego Anderson Dalmolin, Maria João Ramos Pereira, Alexandro Marques Tozetti.

Writing ± review & editing: Diego Anderson Dalmolin, Maria João Ramos Pereira, Alexandro Marques Tozetti.

